# Population Dynamics of Hybrid State during Adaptive Therapy in Cancer

**DOI:** 10.1101/2021.11.27.470219

**Authors:** Ghanendra Singh

## Abstract

Drug resistance emerges due to drug-induced phenotypic switching of drug-sensitive to drug-resistant subpopulations in cancer during therapy. Existing models indicate the competitive advantage of sensitive over resistant population to regulate tumor and reducing the treatment cost with increased time to progression of tumor ultimately benefiting the patient in a clinical setting. Here, we present a Lotka Volterra (LV) based population dynamics (PD) model of the drug-sensitive, drug-resistant, and transient drug-hybrid state along with phenotypic switching during adaptive therapy based on a simple cancer biomarker (CB) to decide the adaptive therapy dosage to regulate cancer. We identified that the strength of intra competition along with phenotypic switching parameters is crucial to mediate the effectiveness of adaptive therapy and also investigated the significance of the initial fraction of subpopulations on AT. We hypothesize and predict the dynamics of drug-induced transient hybrid state playing a key role in the cancer cells undergoing metastasis.

## Introduction

Drug resistance limits the efficacy of therapy for patients with cancer [1]. Researchers are trying to understand the underlying mechanism of drug resistance from molecular, cellular [2], and population dynamics perspectives [3] to solve the problem of drug resistance. Tumor heterogeneity [4] and phenotypic plasticity [5, 6] is present in a population of cancer cells consisting of both drug-sensitive and drug-resistant sub-populations that lead to acquired drug-induced resistance [7] during therapy. Recent studies have indicated the existence of a transient hybrid state during metastasis [8].

Static therapeutic techniques have been less effective due to the tumor microenvironment and phenotypic modification of dynamical cancer systems. Adaptive therapy technique [9] utilizes the competitive advantage of drug-sensitive cells over drug-resistant cells to regulate and delay the reoccurrence of the tumor population as the resistance population pays the cost of fitness compared to the sensitive population. Depending on certain criteria mentioned by [10], AT is effective compared to the conventional maximum tolerated dose therapy (MTD) but if the tumor consists of an aggressive resistant subpopulation, then the effectiveness of AT is comparable to MTD.

Recently, Lotka-Volterra (LV) competition ODE based two models are proposed [11] in which the first model drug-sensitive population inhibits the growth of drug-resistant population competing for resources. In the second model, drug-induced phenotypic switching is also considered. On the same lines, we have considered inherent stochasticity within the heterogeneous population, drug-induced phenotypic plasticity during adaptive therapy, and phenotypic reversibility.

In this paper, we discuss a population dynamics model to understand the competition between drug-sensitive and drug-resistant subpopulations and the role of drug-induced phenotypic switching giving rise to drug resistance during adaptive therapy. Next, to study the role of intra competition among the subpopulation and identify the parameter regimes where phenotypic switching or competition dominates. Later incorporate third drug-induced hybrid phenotype state which gives rise to metastasis to understand the importance of having an intermediate drug-hybrid cell state during adaptive therapy from a population dynamics perspective. Hybrid states indicate high metastasis potential. Does this intermediate or partially resistant or hybrid state present any advantage in terms of phenotypic switching for cancer cells’ decision-making to undergo metastasis during changes in the tumor microenvironment to escape from therapeutic damage?

## Methods

Here we present two LV (Lotka-Voltera) based simple stochastic differential equation models, SR model and SHR model that consider competition between sensitive (S), partially resistant or hybrid (H), and fully resistant (R) populations along with phenotypic switching (PS) and a general cancer biomarker (CB) similar to Prostate Specific Antigen (PSA) in prostrate cancer, Alpha-Fetoprotein (AFP) in liver cancer, BRCA1/2 (breast/ovarian cancer), BRAF/V600E(melanoma/colorectal cancer), and many more. Initially, competition between S and R is considered with PS in the SR model. Later, the role of H is also added in the systems of equations in the SHR model. We also consider two modes of modeling with low and high stochasticity (*σ*). Equations for both models are given below.

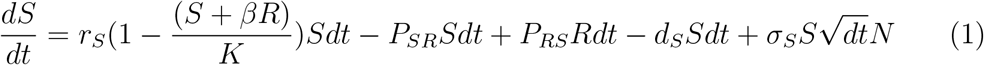

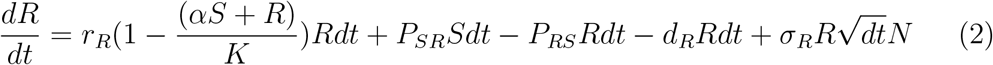

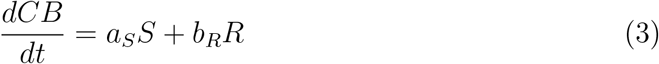

System of equations 1 represents sensitive subpopulation dynamics, 2 represents resistant population dynamics, and 3 represents cancer biomarker dynamics to calculate AT dosage criteria for the SR model. Similarly system of equations 4, 5, 6, and 7 represents sensitive, hybrid, resistant, cancer biomarker dynamics for the SHR model respectively. Definitions for the equation parameters and values are given in Table 1.

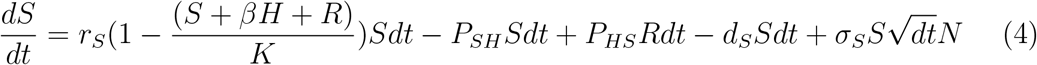

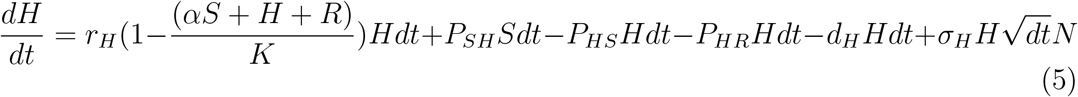

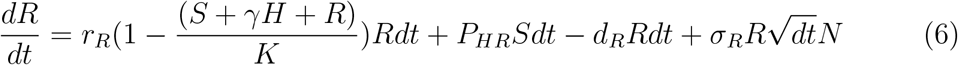

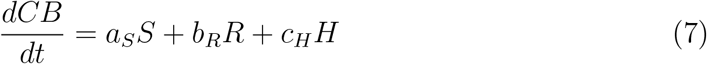

**Table 1:** Model parameters

| Parameters | Description | Values |
| --- | --- | --- |
| $r_S, r_H, r_R$ | Growth rates for S,H, and R | 0.2 |
| $d_S, d_H, d_R$ | Death rates for S,H, and R | 0.1 |
| $P_{SR}, P_{RS}$ | SR model switching rates | 0 - 1 |
| $P_{SH}, P_{HS}, P_{HR}$ | SHR model switching rates | 0 - 1 |
| $\alpha, \beta, \gamma$ | Competition parameters | 0 - 2 |
| $\sigma_S, \sigma_H, \sigma_R$ | Stochasticity of model | 0.001 |
| $a_S, b_R, c_H$ | Cancer biomarker coefficients | 0.6, 0.3, 0.1 |
| $N(0)$ | Initial population of S,H, and R | $10^4$ |
| $K$ | Carrying capacity | $10^5$ |
| $F_R$ | Initial fraction R(0) at t=0 | 0.01 |
| $S_c$ | CB Critical threshold | 20000 |
| $dt$ | Time step | 1 |
| N | Normal distribution $\mu=0, \sigma=1$ | $N(0,1)$ |

## Results

### Competition without Phenotypic Switching in the SR Model

Simulation result of the SR model and its temporal population dynamics. In figure 3a, when initial resistant population R(0) fraction is low *F_R_* =1% at time t=0, it takes large time for resistant population to recover. In figure 3b, if the initial fraction R(0) is medium *F_R_*=10%, it takes lesser time compared to the previous case, and in figure 3c, if the initial fraction of resistant population R(0) is equal to the initial population of S(0), R quickly takes over. It indicates the importance of having an initial low resistant population meeting the prerequisites mentioned by [10] for the effectiveness of AT without PS.

**Figure 1:**
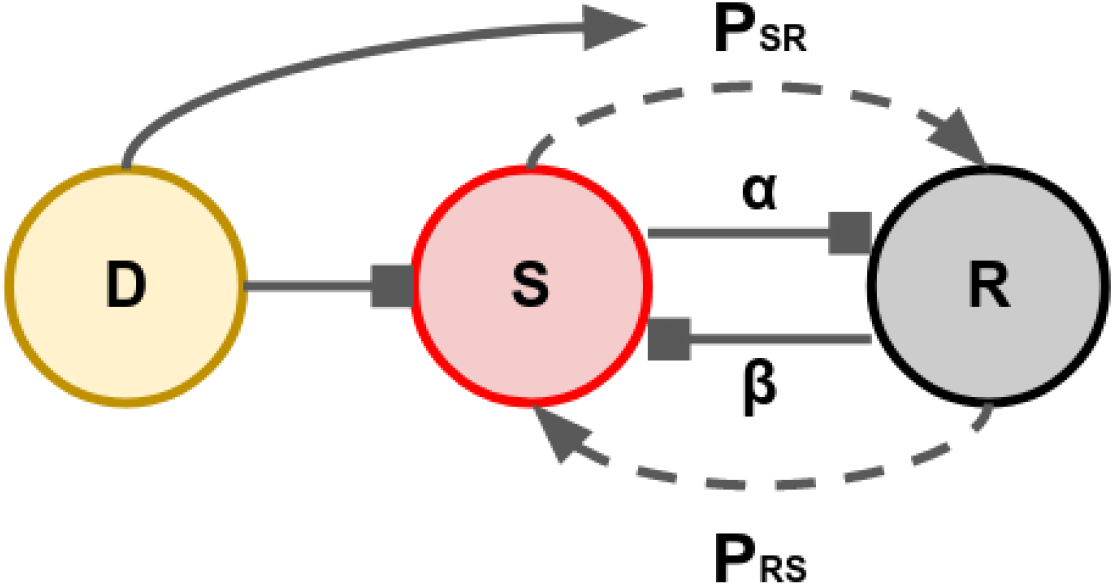
Sensitive and Resistant (SR) population dynamics model

**Figure 2:**
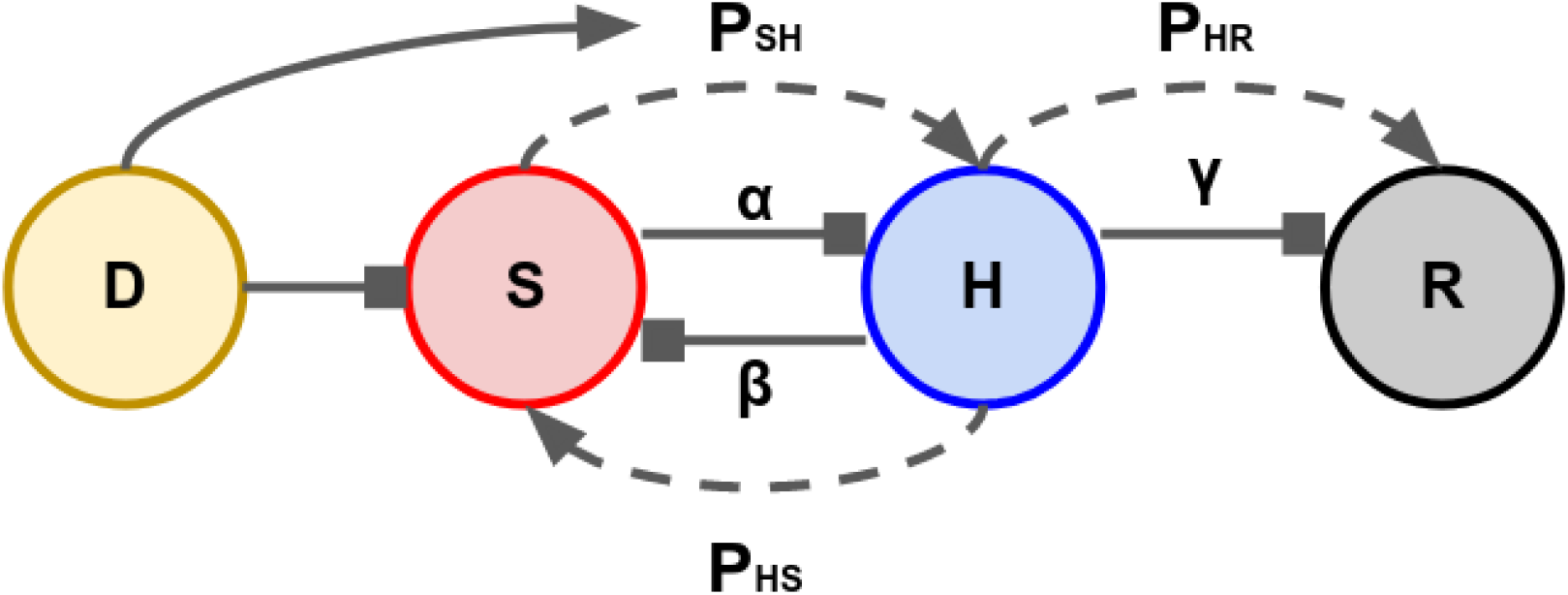
Sensitive, Hybrid, and Resistant (SHR) population dynamics model

**Figure 3:**
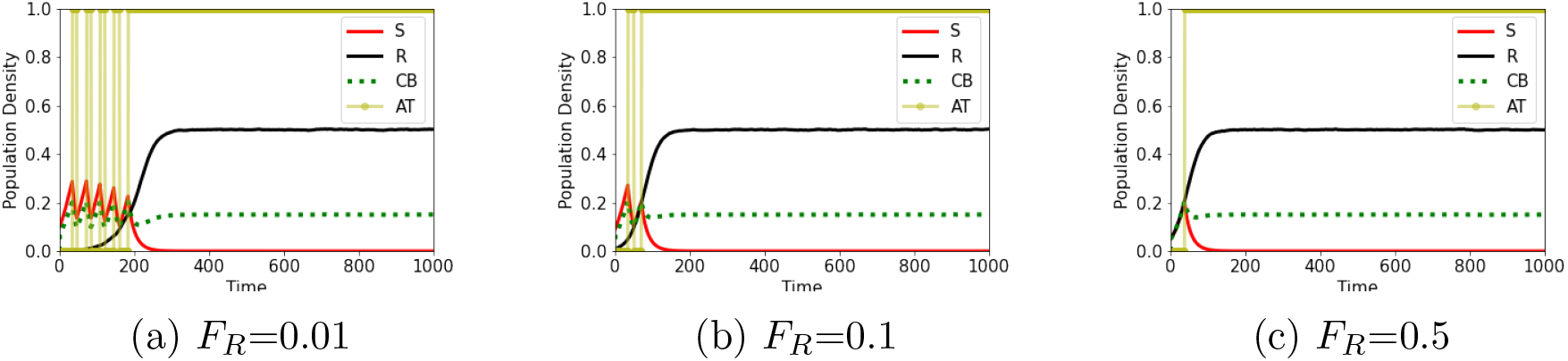
Role of initial fraction *F_R_* in competition (C) between S and R without PS. (a) Initial resistant population R(0) *F_R_* = 0.01. (b) *α* = 1.0 and Fr: 0.1 (c) *F_R_*=0.5

### Competition with Phenotypic Switching in the SR Model

Now, phenotypic switching (PS) *P_SR_*=0.01 is introduced along with pheotypic reversibility *P_RS_*=0.02, and the simulation results are shown in figure 4 for different initial fractions FR. In figure 4a, initially, when there is no drug induced PS from S to R, occurance of resistance population is delayed and when PS is introduced, the resistant population quickly takes over. Similarly in figures 4d and 4f with different initial fractions, resistant population takes over quickly.

**Figure 4:**
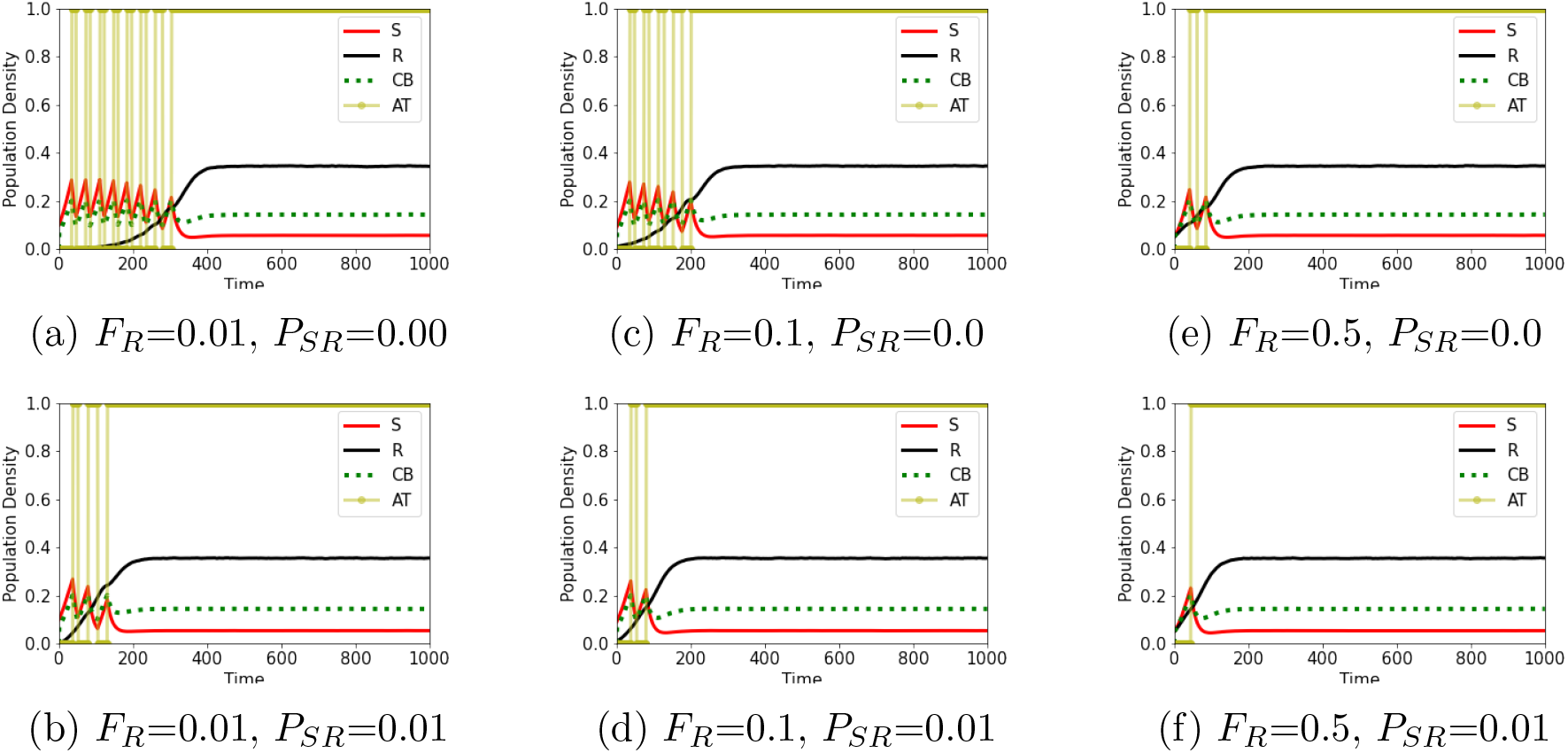
Competition(C) between S and R with PS. *α* = 1.0 and *β* = 0.5, *P_SR_* = 0.0 or 0.01 and *P_RS_* = 0.02. With gradual increase in the initial fraction *F_R_* of resistant population R(t=0), effectiveness of the AT reduces.

### Competition and Phenotypic Switching in the SHR Model

#### Default SHR Model

Addition of an intermediate transient drug-hybrid state between drug-sensitive and drug-resistant subpopulation in the SR model gives SHR model. Simulation results are shown in the figure 5 with low stochasticity *σ* and high stochasticity *σ* in figure 5b consisting of all different sub type populations during adaptive therapy regulating the sensitive population without any competition and without phenotypic switching between the subpopulation in tumor. It is referred as default version of SHR model.

**Figure 5:**
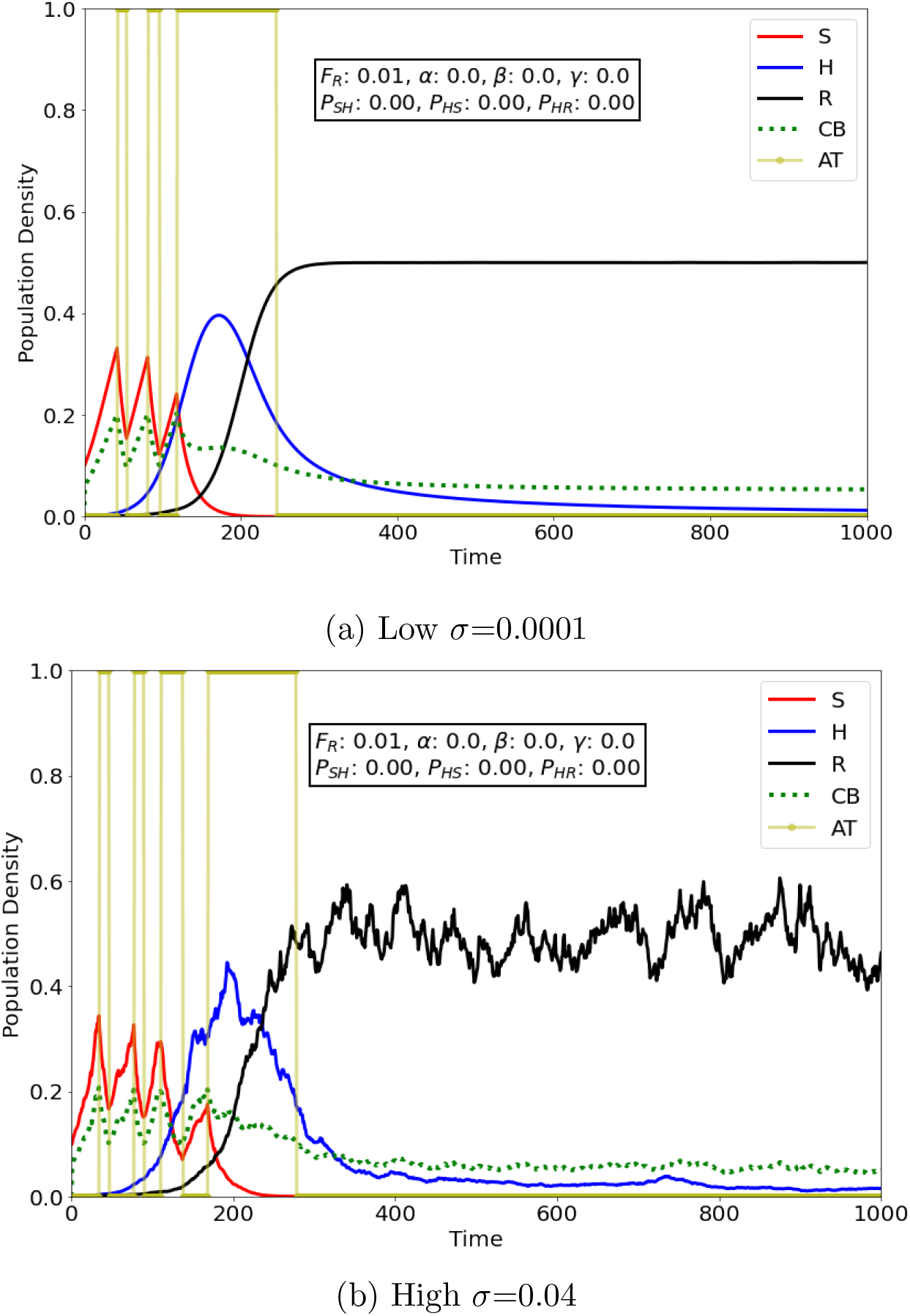
Population dynamics of drug-sensitive (S), drug-hybrid (H) and drugresistance (R) cell states in tumor during adaptive therapy (AT) based on cancer biomarker (CB) without competition *α*=0, *β*=0, *γ*=0 and without phenotypic switching (PS) *P_SH_*=*P_HS_*=*P_HR_*=0. Hybrid cell state exists when S population is reduced due to AT and R population is low. Due to death rate term, population density (PD) of R remains below 1. Parameters for the simulation results are present in Table 1.

#### Transient Hybrid States

To understand the role of hybrid state, we varied and tuned the model parameters and observed interesting behavior. In figure 7a with medium competition parameter *β* value = 0.5 and low H to R PS *P_HR_*=0.001, initially AT is on when high CB threshold is crossed, S population is reduced as general, assuming H cells don’t contribute equally in rising the CB levels up, low CB threshold is not reached and AT is on during which H populations increases. As soon as CB reduces to half of its previous value, AT is off and S cells starts growing and cycle repeats. Similar effect is observed in 7b having low *β*=0.1 but with quick cycles of AT indicating low doses in small interval. In 7c with low competition between S and H and nominal H to R PS *P_HR_*=0.01, we require low AT doses at larger intervals to regulate tumor which is more effective than previous case.

**Figure 6:**
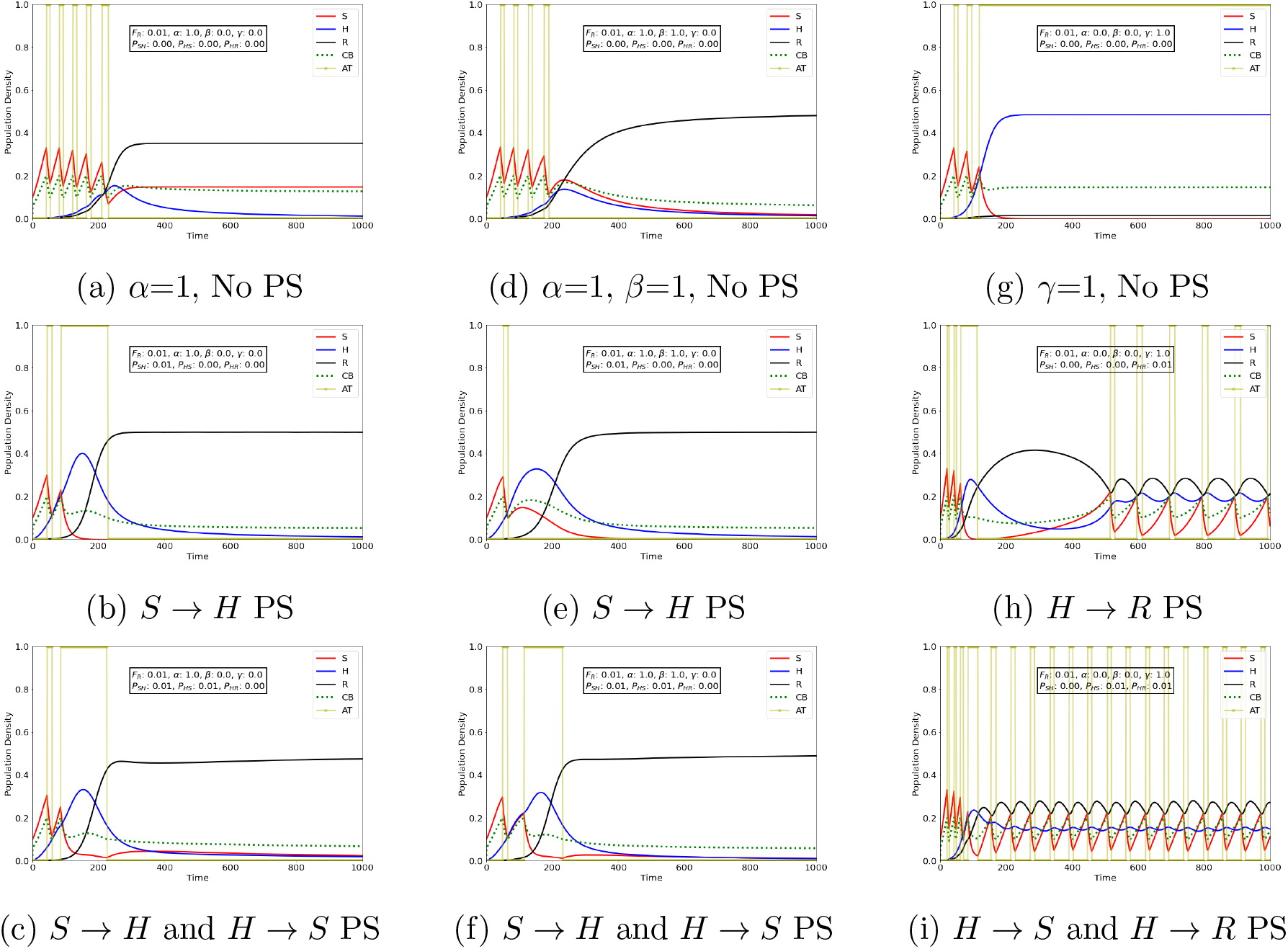
Competition between S, H, and R with/without PS. (*a*) S competes with H. (*b*) S competes with H and AT induced PS from S to H. (*c*) S and H both competes with each other. AT induced PS from S to H and H to S PS due to PR. (*d*) S and H both compete with each other. (*e*) S and H competition along with AT induces S to H PS. (*f*) S and H competition with AT induces S to H PS and H to S PS due to PR. (*g*) H competes with R and takes over at steady state. (*h*) H competes with R along with PS from H to R resulting in a delayed AT based oscillations. (*i*) H competes with R and PS of H to S and R giving rise to an AT based oscillatory behavior regulating R population.

**Figure 7:**
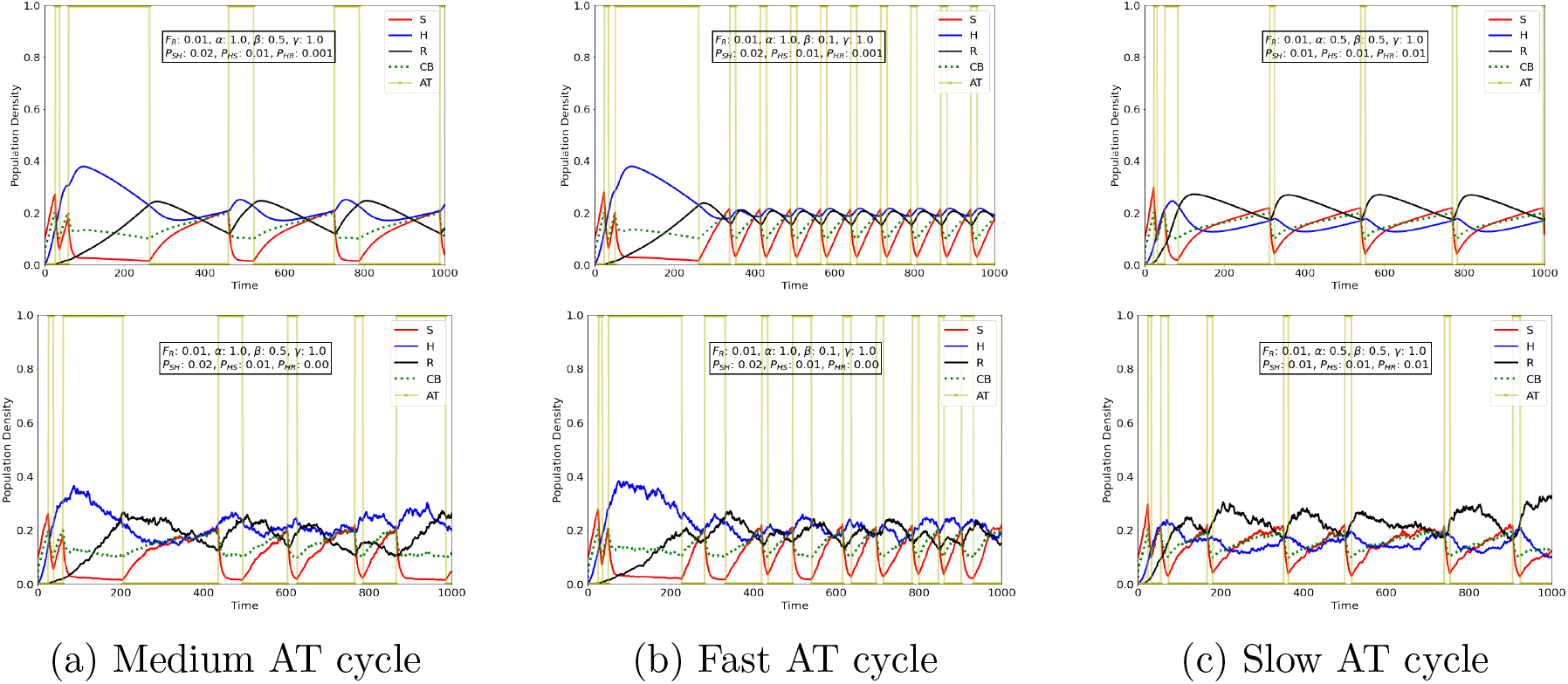
Hybrid state transition between S and R subpopulation. Above figures with low *σ*=0.001 and below figures with high *σ*=0.02. More no. of AT dosage cycles are needed to regulate the inherent stochasticity in the population.

## Conclusions

In this research, we developed a Lotka-Voltera (LV) based two simple mathematical SR and SHR models to understand the population dynamics of drug-sensitive, drug-hybrid, and drug-resistant tumor sub population and used a simple cancer biomarker (CB) threshold for adaptive therapy using stochastic differential equation. We identified certain key parameter regimes where AT is effective and tried to phenomenological explain the importance of intermediate drug-hybrid state role in cancer.

## Future Work

In future, we plan to include the role of genetic mutations leading to a higher fitness of the resistance cells compared to sensitive cells and the role of hybrid state in giving rise to metastatic population with stemness properties. Later, to include the role of growth factors secreted in the tumormicroenvironment (TME) and connect steady state behavior of the cellular mechanisms with population dynamics. Also add drug dynamics to model dose response curve to study its effect and identify minimum and maximum concentration of drugs for high efficacy during treatment.

## Supplementary Material

Below table 1 contains all the parameters and its values used in the SR and SHR model. A threshold value *S_c_*=20000 of Sensitive (S) population is used to calculate cancer biomarker (CB) for adaptive therapy (AT). If CB is higher than *S_c_* then AT in turned on during which growth rate *r_S_* of sensitive population is modified to 0.1 and death rate to *d_S_*=0.2. When CB reached half of its initial value, AT is turned off, parameters *r_S_*=0.2 and *d_S_*=0.1 takes default value. No changes are made in the growth and death rates for hybrid and resistant population for these models.

## Notes

https://github.com/Ghanendra19213/Adaptive-Therapy/blob/main/PD%20of%20Hybrid%20State%20during%20AT%20in%20Cancer.ipynb

### Competing Interest Statement

The authors have declared no competing interest.

